# Cross-domain information fusion for enhanced cell population delineation in single-cell spatial-omics data

**DOI:** 10.1101/2024.05.12.593710

**Authors:** Bokai Zhu, Sheng Gao, Shuxiao Chen, Jason Yeung, Yunhao Bai, Amy Y. Huang, Yao Yu Yeo, Guanrui Liao, Shulin Mao, Sizun Jiang, Scott J. Rodig, Alex K. Shalek, Garry P. Nolan, Sizun Jiang, Zongming Ma

**Author notes:** Equal Contributions (Bokai Zhu, Sheng Gao, Shuxiao Chen). Senior Authors (Sizun Jiang, Garry P. Nolan, Sizun Jiang, Zongming Ma).

## Abstract

Cell population delineation and identification is an essential step in single-cell and spatial-omics studies. Spatial-omics technologies can simultaneously measure information from three complementary domains related to this task: expression levels of a panel of molecular biomarkers at single-cell resolution, relative positions of cells, and images of tissue sections, but existing computational methods for performing this task on single-cell spatial-omics datasets often relinquish information from one or more domains. The additional reliance on the availability of “atlas” training or reference datasets limits cell type discovery to well-defined but limited cell population labels, thus posing major challenges for using these methods in practice. Successful integration of all three domains presents an opportunity for uncovering cell populations that are functionally stratified by their spatial contexts at cellular and tissue levels: the key motivation for employing spatial-omics technologies in the first place.

In this work, we introduce <u>Cell S</u>patio- and <u>N</u>eighborhood-informed <u>A</u>nnotation and <u>P</u>atterning (CellSNAP), a self-supervised computational method that learns a representation vector for each cell in tissue samples measured by spatial-omics technologies at the single-cell or finer resolution. The learned representation vector fuses information about the corresponding cell across all three aforementioned domains. By applying CellSNAP to datasets spanning both spatial proteomic and spatial transcriptomic modalities, and across different tissue types and disease settings, we show that CellSNAP markedly enhances *de novo* discovery of biologically relevant cell populations at fine granularity, beyond current approaches, by fully integrating cells’ molecular profiles with cellular neighborhood and tissue image information.

## Introduction

There has been a recent surge in the development of multiplexed imaging technologies capable of simultaneously evaluating 40-100 protein targets (1–3) or several hundred to thousands of mRNA targets (4–6) at single-cell or sub-cellular resolution, within their native tissue context. High-plex *in situ* imaging has facilitated the discovery of intricate tissue structures and local neighborhoods, paving the way for novel approaches to disease interference (7–9). These high-dimensional “spatial-omics” data often require sophisticated approaches for data analysis to uncover biologically meaningful insights. In what follows, we refer to spatial-omics data at single-cell or finer resolution as single-cell spatial-omics data. The majority of information in these data is contained in the following three complementary domains (10): 1) single-cell expression levels of measured biomarkers, 2) relative locations of cells, and 3) image of measured tissues, providing information related to cellular morphology and tissue architecture, through multiple channels. For simplicity, we refer to them here and after as 1) expression, 2) location, and 3) image domains, respectively.

Analysis of single-cell spatial-omics data begins with cell phenotyping: Single-cell boundary masks are generated from raw images by cell segmentation approaches (e.g., (11, 12)), which are then employed to extract the expression level of molecular features from each cell to create a cell-by-feature data matrix. The data matrix is often treated as from a dissociated single-cell study for cell population identification. In the most bare-bone form, the process involves applying a clustering algorithm (e.g., (13–16)) on the matrix, followed by cell type annotation by a human expert who compares highly-expressed biomarkers of each cluster to known markers of a list of pre-defined cell populations. Conventionally, the foregoing cell population delineation (i.e., clustering cells into groups with distinct bio-molecular profiles and/or biological functionalities) and identification (i.e., cell type annotation) process rely solely on feature expression levels, whilst image-level information (e.g., cell relative location, marker/tissue image information) is only utilized during downstream analyses, such as cellular neighborhood or tissue schematic identification (17–20). As high spatial resolution is often accompanied by a limited targeted panel size, this (molecular-)feature-only approach only distinguishes coarser cell populations when compared with high-throughput dissociated single-cell data, thus severely hindering the realization of the full potential of spatial-omics studies.

Tissue image information at different scales can facilitate invaluable insights into cellular-level processes, including but not limited to cell type and activation state. This is exemplified by how pathologists are able to identify certain cell populations from H&E images, based on information contained in cell morphology, location, and surrounding tissue architecture (21, 22). In addition, spatial information in an image can further differentiate subpopulations of cells, which cannot be fully captured by the cell’s measured features alone, but rather in concert with the distinct microenvironments that drive cell states beyond cell identities. Furthermore, the image-level information can complement the molecular profile information confined by the antibody or RNA probe panel size. This is exemplified by the distinctive effector and suppressor activities of T cells in close contact with tumors, compared to those farther away (23–25). In parallel, B cells located in different layers of niches within the germinal center also represent different populations with distinct functionality and cell state changes (26, 27). Therefore, ideal cell population delineation and identification in spatial-omics data should take advantage of spatial and tissue image information availability.

When one has a set of pre-defined cell populations in the form of an annotated training dataset, new methods (28, 29) have been proposed to leverage the relative locations of cells in a spatial-omics dataset when classifying the cells to the pre-specified populations. However, as classification methods, they are supervised-learning by nature, and hence do not directly accommodate the definition and discovery of new cell populations unseen in the training data or the partitioning of a population into multiple biologically distinct subpopulations. In addition, image domain information is ignored. For cell population delineation, recent methodologies have explored fusing single-cell morphology (i.e., individual cell shape information) (30) and spot spatial adjacency information (i.e., a form of relative location information among cells) (31) to the original expression profiles, for generating new numerical representations of cells that could better represent their differences than the measured molecular profiles. Compared with classification, the primary advantage of applying unsupervised clustering methods on these cell representation vectors is the possibility of *de novo* cell population discovery: Clustering methods make no assumption about the existing cell populations in data, making the identification of previously unseen biological events more accessible.

In view of the gap between the available information in data and that leveraged by state-of-the-art computational methods, we postulated that integrating information contained in the location and the image domains of spatialomics data with single-cell molecular profiles would notably enhance the granularity when delineating cell populations. We thus present CellSNAP (Cell Spatio- and Neighborhood-informed Annotation and Patterning), an unsupervised information fusion algorithm, broadly applicable to different single-cell spatial-omics data modalities, for learning cross-domain integrative single-cell representation vectors. In particular, CellSNAP-learned representation vectors incorporate information from expression, location, and image domains, hence exhausting major information sources in single-cell spatial-omics data. Existing unsupervised clustering algorithms, such as Leiden clustering and its peers, can operate directly on the CellSNAP-learned representations rather than the conventional feature expressions, enabling spatio-informed fine-grained delineation and *de novo* discovery of cell populations across diverse imaging modalities and biological samples.

## Results

### Overview of CellSNAP

The following is a brief description of the CellSNAP pipeline (**Fig. 1**). See Material and Methods for more technical details. CellSNAP is an information fusion algorithm operating on single-cell spatialomics modalities, including but not necessarily limited to both spatial proteomics (e.g., CODEX (3)) and spatial transcriptomics (e.g., CosMx-SMI (6)). To start with, CellSNAP curates three pieces of inputs *for each cell* corresponding to the three information domains: 1) expression domain: an expression profile vector, e.g., a vector of protein expressions from a CODEX experiment; 2) location domain: a cellular neighborhood context vector, i.e., a vector recording composition information of neighboring cells around each cell; 3) image domain: an image tensor recording local tissue images in multiple channels i.e., a collection of images that capture each cell’s adjacent and local tissue image patterns. To access the neighborhood context information, we assume that each cell has been given a coarse initial cluster label. Hence, its neighborhood context can be represented by the proportions of its spatial nearest neighbors in different initial clusters. When *a priori* human expert annotation is not available, one can obtain initial cluster labels by grouping the cells according to their expression profile similarities. When human-expert cell type annotations are available, the annotated labels can serve as initial cluster labels. For all results reported in this paper, we used Leiden clustering (16) with resolution level 0.5 on a feature-expression induced nearest neighbor graph of each dataset for generating initial cluster labels, *without* human intervention.

**Figure 1:**
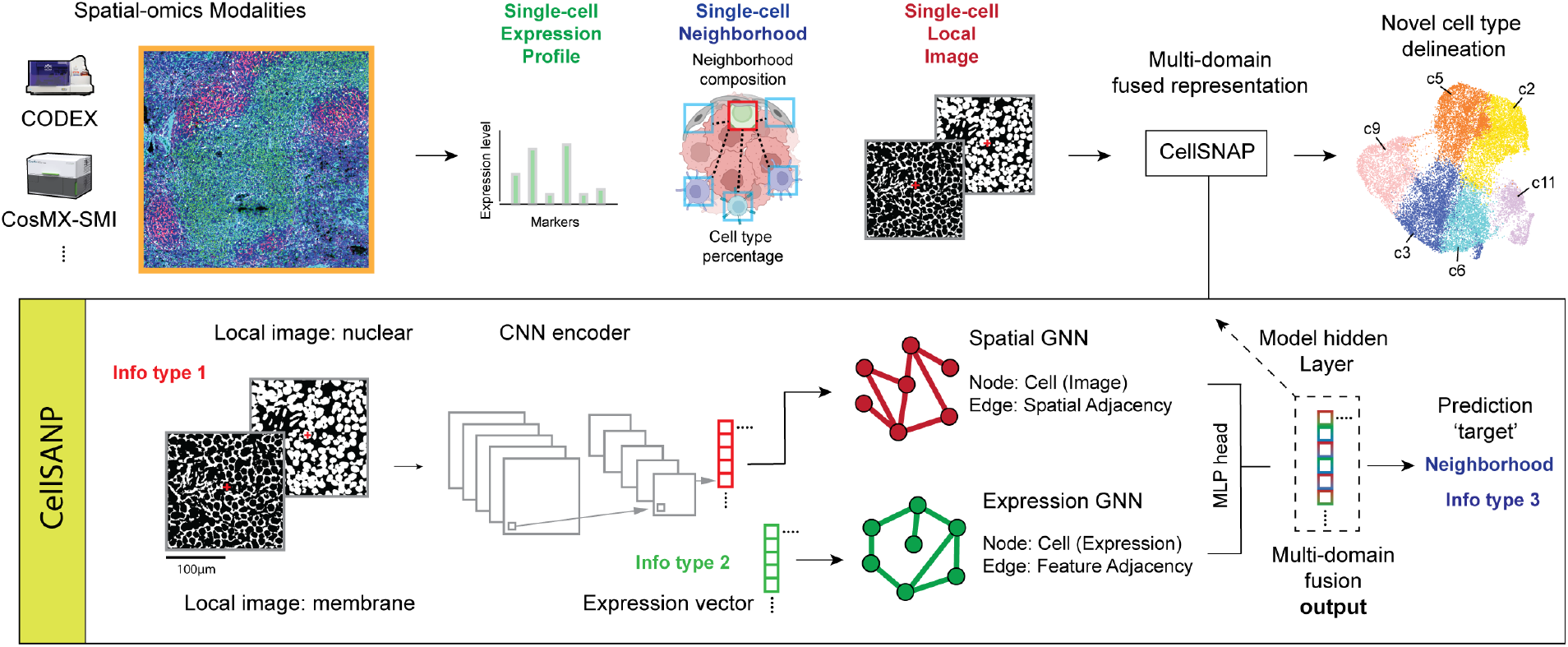
Illustration of the CellSNAP pipeline. CellSNAP is compatible with imaging-based spatial-omics modalities with single-cell or finer resolutions (e.g., CODEX and CosMx). Information from three domains is extracted from each individual cell and its surroundings: 1) single-cell expression profile (e.g., measured protein or mRNA features); 2) single-cell location information (e.g., cellular neighborhood composition); 3) single-cell local tissue image information (e.g., local images from nuclear and membrane channels). CellSNAP takes these three types of information as input. It first utilizes a CNN encoder to extract features from images of local tissues surrounding each cell. Next, two separate GNN models are constructed in parallel: 1) a ‘Spatial-GNN’, where each node represents a cell, with the initial node vector assigned as CNN-extracted local image features, and nodes are connected according to spatial adjacency. 2) an ‘Expression-GNN’, where each node represents a cell, with the initial node vector assigned as the expression profile, and nodes are connected according to expression similarity. The two GNNs are connected by an overarching MLP head which combines the message passing outputs of the two GNNs for predicting the target vector of each cell, that is the concatenation of the cell’s feature-based population identity (one-hot) and its neighborhood composition (percentage) vectors. After training, the last layers of the two GNN models are extracted, combined, and reduced (via SVD) to form the final, tri-domain integrated representation vector for each cell. This multi-domain fused representation vector is then used in downstream analysis for cell type identification purposes, which is compatible with commonly used unsupervised clustering methods (e.g., Leiden clustering). Detailed illustration of the involved model architectures can be found in **Supp. Fig. 1**.

CellSNAP leverages a novel neural network architecture, which we term SNAP-GNN-duo (**Supp. Fig. 1**), for orchestrated information integration across three domains. The architecture consists of two parallel Graph Neural Networks (GNNs) (32) with identical node set but distinct network topology, connected by an overarching multi-layer perceptron (MLP) (33) head. In both GNNs, nodes represent cells in the measured tissue section(s) with one-to-one correspondence. One GNN (the Spatial GNN in **Fig. 1**) is constructed on a spatial adjacency graph, where for each node its nodal feature is initialized with the local tissue image encoding vector of the cell (see next paragraph for details), and pairs of nodes are connected if their spatial locations in the tissue section are close. The other GNN (the Expression GNN in **Fig. 1**) is built on a feature similarity graph, where for each node its nodal feature is initialized with the cell’s expression profile, and pairs of cells are connected if their expression profiles are similar.

To compute the local tissue image encoding vector for each cell, we train from scratch a Convolutional Neural Network (CNN) model (34), which we call SNAP-CNN (**Supp. Fig. 1**), as an image encoder. SNAP-CNN takes each cell’s local tissue image, processes it through an AlexNet-like architecture (34), and predicts the cell’s cellular neighborhood context vector. The resulting fitted encoding vectors from training (a pre-specified hidden layer in the trained SNAP-CNN) are used as local tissue image encoding vectors for individual cells. SNAP-CNN supplies values for initializing SNAP-GNN-duo, and its training (details in Material and Methods) is performed prior to that of SNAP-GNN-duo.

After initialization and parallel message passing on the SNAP-GNN-duo, the updated nodal vectors of the two GNNs are concatenated and used as inputs for the final MLP head. The MLP head trains and predicts for each cell the target vector which is the concatenation of the cell’s cellular neighborhood context vector and a one-hot vector recording the cell’s initial cluster label. Finally, after training, a designated hidden layer of the MLP head is used as the output representation vector of the CellSNAP pipeline (**Supp. Fig. 1**). See Material and Methods for details on training SNAP-GNN-duo.

By design, the CellSNAP representation fuses single-cell expression, cellular neighborhood, and local tissue image information. In downstream analysis, an existing clustering algorithm (13–16) can be directly applied to the CellSNAP representation vectors for cell population delineation at fine granularity, with no further modification required. For benchmarking purposes, we applied Leiden clustering (16) on the resulting CellSNAP representation in this study to showcase its applications. Human experts and/or machine learning algorithms can then determine the biological states of the CellSNAP clusters and perform further downstream investigations.

### Application to CODEX lymphoid tissue data

We performed CellSNAP on a healthy mouse spleen CODEX dataset (3), which includes 30 protein markers and 53,500 cells. To quantitatively evaluate the capacity to delineate cell populations of various cell representation methods, we implemented four metrics: Silhouette Score, Calinski-Harabasz Index, Davies-Bouldin Index, and Modularity Score (see Material and Methods for details). For These metrics do not require ground truth cell population information, and thus are suitable for objectively benchmarking the performance of different methods for *de novo* cell type identification. We computed these metrics based on single-cell representation vectors obtained from five different methods, including: 1) Feature: each cell’s feature expression vector; 2) Concatenation: the concatenation of each cell’s feature expression vector and neighborhood composition vector; 3) SpiceMix (31): each cell’s SpiceMix representation vector; 4) MUSE (30): each cell’s MUSE representation vector; 5) CellSNAP: each cell’s CellSNAP representation vector. We observed improved clustering performance with the CellSNAP representation, compared to other methods on the mouse spleen CODEX data (**Fig. 2A**), as defined by the metrics above.

**Figure 2:**
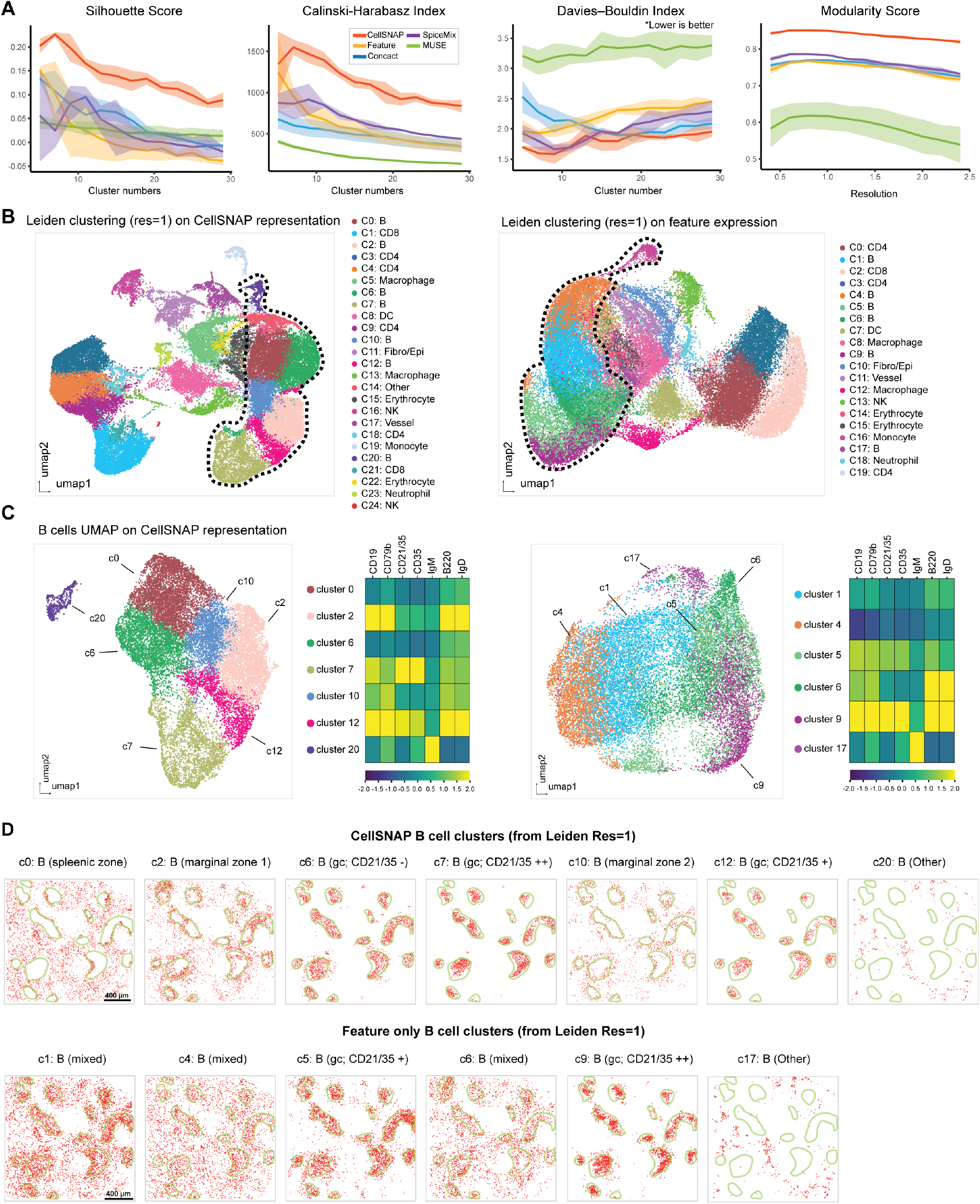
Refined B cell subpopulations discovered by CellSNAP in a healthy mouse spleen CODEX dataset. **(A)** Metric-based evaluations of cell population delineation performances on CODEX mouse spleen tissue. Representations of cells, from 5 different methods, were used as input: CellSNP representation, feature (protein expression table), concact (protein expression + neighborhood composition table), SpiceMix representation, and MUSE representation (detail in Material & Methods). A total of 5 batches, each with 10,000 randomly selected cells were tested. Solid line indicates average and shade indicates 95% CI of the scores. **(B)** UMAP visualizations of representations and Leiden clustering results. Cell types of the CellSNAP or feature-only clusters were annotated based on the average expression profiles of the clusters. Left panel: CellSNAP representation; Right panel: feature expression. Dotted line indicates B cell subpopulations. **(C)** UMAP visualizations and B cell-related protein expression profiles of clusters annotated as B cells. Left panel: B cell clusters from CellSNAP and their expression heatmap. Right panel: B cell clusters from feature expression and their expression heatmap. **(D)** Comparison of spatial locations of different B cell clusters identified by CellSNAP representation vs. feature expression in the spleen tissue. In each plot, red dots indicate cells from a specific cluster, and green lines indicate germinal center boundaries. Upper panel: Spatial locations of B cell subpopulations identified by CellSNAP representation clusters. Lower panel: Spatial locations of B cell subpopulations identified by feature expression clusters.

We next performed a more nuanced comparison of cell type identifications based on clustering results (from Leiden clustering with resolution = 1) on: 1) CellSNAP representation, or 2) feature-only expression profile (i.e., expressions of 30 protein markers). First coarse cell type annotations were generated based on the protein profiles from either CellSNAP or feature-only clusters (**Fig. 2B, Supp. Fig. 2**). Subsequently, we focused on clusters annotated as B cells. B cells were chosen as they encompassed the largest population in this dataset (*≈* 45% of all cells). The CellSNAP and feature expression representations produced 7 and 6 Leiden clusters which were annotated as B cells respectively (**Fig. 2C**). B cell clusters from the feature expression representation were more intermixed when visualized via UMAP, and less distinctive in regards to B cell-related protein marker expression (**Fig. 2C, right**); B cell clusters from the CellSNAP representation were more well-structured when visualized via UMAP, and more distinctive in terms of B cell-related protein marker expression (**Fig. 2C, left**). To further validate our findings that B cell subpopulations were better stratified using CellSNAP, we investigated the biological relevance of the identified B cell clusters by observing their spatial localization (**Fig. 2D**). While the conventional features-only approach could partially delineate B cell subpopulations, including the identification of germinal center (GC) B cells with medium or high CD21/CD35 expression, most Leiden clusters from feature representation (c1, c4, c6) were mixtures of different B cell subgroups (**Fig. 2D, bottom**). In comparison, Leiden clusters from CellSNAP representation spatially delineated the B cell subgroups successfully, including not only the GC B cells with medium or high CD21/CD35 expression as identified in the feature-only approach, but also GC CD21/CD35 negative B cell (c6), spleenic zone B cell (c1), and marginal zone B cell subgroups (c1, c10) (**Fig. 2D, top**) (35–37).

We further evaluated CellSNAP for its cell population delineation performance on a human tonsil CODEX data (38), consisting of 46 protein markers across 102,574 cells. Akin to the mouse spleen data, we observed improved clustering performance based on the quantitative benchmarking metrics (**Supp. Fig 3A**). We similarly performed cell type annotation on both CellSNAP representation and feature-only representation (**Supp. Fig. 3B, Supp. Fig. 4**), and identified non-overlapping B cell annotations between CellSNAP and feature-only results: whilst both methods robustly uncovered general B cells (including GC B cells), CellSNAP was able to identify an additional population that was not identified with the feature-only representation (cells labeled as red, **Supp. Fig. 3C**). This replicating non-GC B cell population was identified using CellSNAP as a distinctive Leiden cluster (CellSNAP - c10 in **Supp. Fig. 3B**), but was intermixed with other B cells in a feature-only Leiden cluster (feature - c8 in **Supp. Fig. 3B**). The failure of the feature-only approach in distinguishing this B cell subpopulation is likely due to its similar protein expression profile to other GC B cells, both exhibiting high levels of the proliferation marker Ki67 in CellSNAP clusters c8 and c10 (**Supp. Fig. 3E**). A close visual inspection of the spatial localization of CellSNAP - c10 cluster cells confirmed their distinctive arrangements bordering GCs (**Supp. Fig. 3F**).

### Application to HCC cancer tissue CosMx-SMI data

Given the compelling evaluation of CellSNAP on spatial proteomics data, we further extended its applicability on cell population delineation with spatial transcriptomics data. We first deployed CellSNAP and other methods on a Hepatocellular carcinoma (HCC) tumor CosMx-SMI dataset (48). This dataset consists of 54,867 cells, with an RNA panel of 997 genes. We first evaluated the cell clustering performance using the previously described quantitative metrics, and continued to observe superior performance from CellSNAP (**Fig. 4A**). We then performed cell type annotation based on Leiden clustering results, either from CellSNAP representation or feature-only representation (**Supp. Fig. 6A, B**). Between the two methods, CellSNAP allowed finer delineation of cell subpopulations. For instance, 2 clusters of macrophages were identified using feature-only representation (**Supp. Fig. 6A**,**B**). In contrast, an increased number of macrophage clusters, each with a distinctive niche occupation, were obtained from CellSNAP representation (**Fig. 4B**). We observed that the macrophage subpopulation CellSNAP c6 exhibited a unique spatial occupation of the Tumor-Immune interface, which was missing in the feature-only results (**Fig. 4B, Supp. Fig. 6C**). Further investigation consolidated the functional differences of subpopulation CellSNAP - c6 from other macrophages in this dataset. Differential expression (DE) analysis between c6 macrophages and other macrophages revealed upregulated genes unique for CellSNAP - c6, including the *C1Q* family (*C1QA, C1QB, C1QC*), *LYZ*, and the *HLA* family (*HLA*.*DQA1, HLA*.*DPA1*) (**Fig. 4C**).

**Figure 3:**
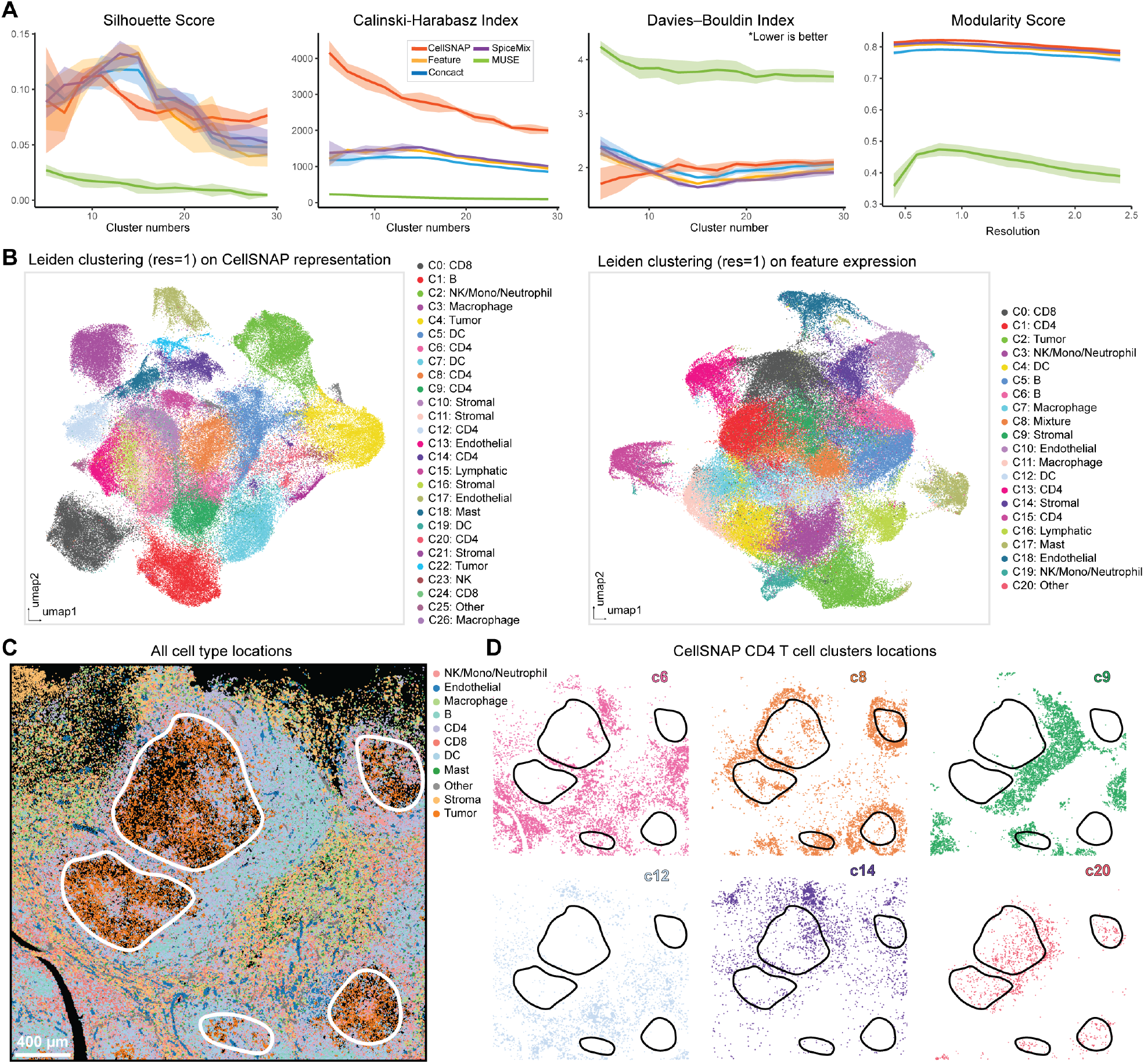
Refined T cell subpopulations in tumor microenvironments discovered by CellSNAP in a cHL tumor CODEX dataset. **(A)** Metric-based evaluations of cell population delineation performances on CODEX human cHL tissue. Representations of cells, from 5 different methods, were used as input: CellSNP representation, feature (protein expression table), concact (protein expression + neighborhood composition table), SpiceMix representation, and MUSE representation (detail in Material & Methods). A total of 5 batches, each with 10,000 randomly selected cells were tested. Solid line indicates average and shade indicates 95% CI of the scores. **(B)** UMAP visualizations of representations and Leiden clustering results. Cell types of the CellSNAP or the feature-only clusters were annotated based on the average expression profiles of the clusters. Left panel: CellSNAP representation; Right panel: feature expression. **(C)** Visualization of cell type spatial locations in the cHL tissue, colored by annotations on CellSNAP clusters. Black regions are empty spaces. White lines indicate the borders of the cHL tumor regions. **(D)** Visualization of the spatial locations of different CD4 T cell subpopulations identified by CellSNAP representation clusters. Black lines indicate borders of the cHL tumor regions.

**Figure 4:**
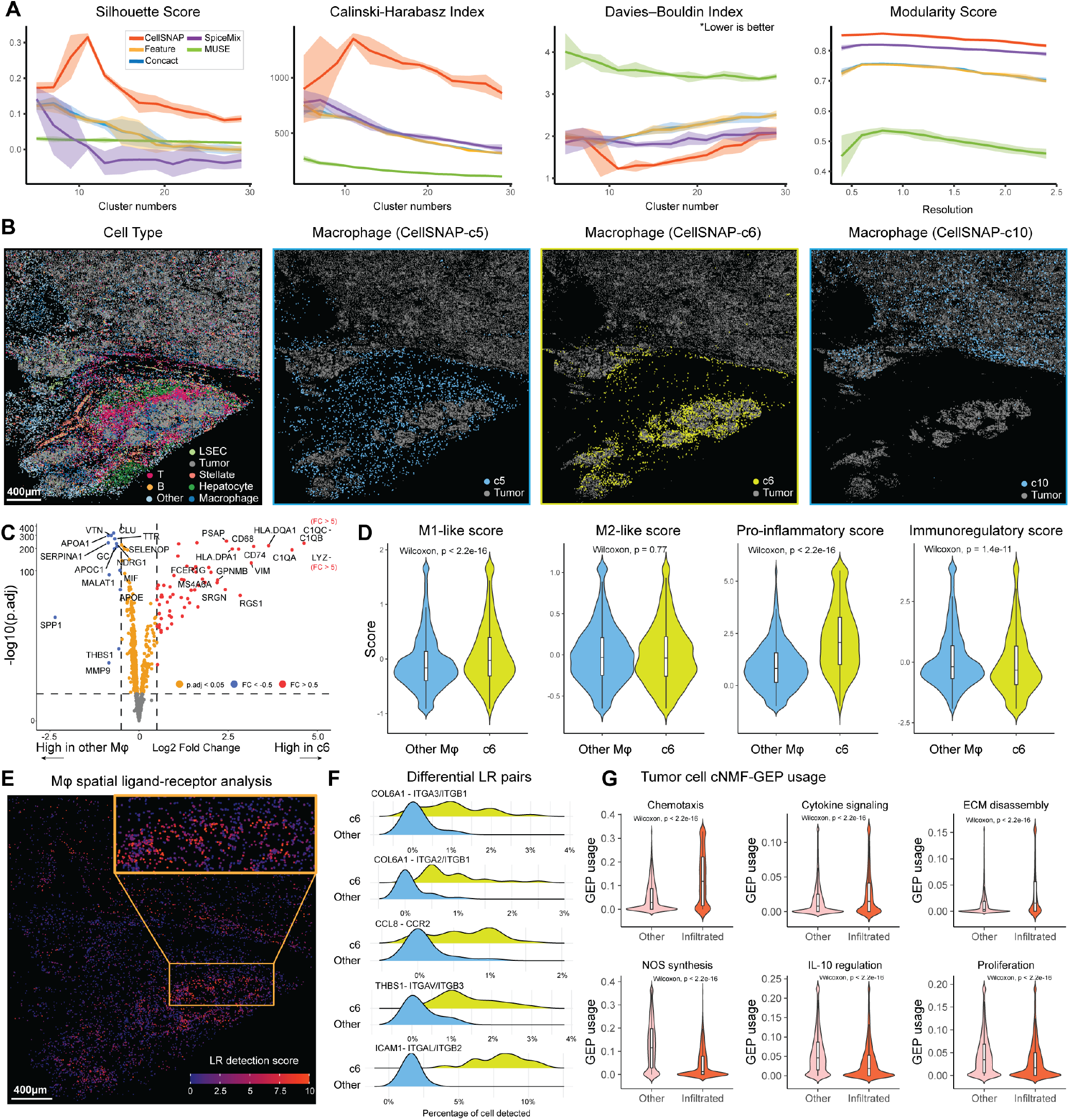
CellSNAP-enabled delineation of biologically distinct macrophage subpopulations in a HCC tumor CosMx-SMI dataset. **(A)** Metric-based evaluations of cell population delineation performances on CosMx-SMI human HCC tissue. Representations of cells, from 5 different methods, were used as input: CellSNP representation, feature (protein expression table), concact (protein expression + neighborhood composition table), SpiceMix representation, and MUSE representation (detail in Material & Methods). A total of 5 batches, each with 10,000 randomly selected cells were tested. Solid line indicates average and shade indicates 95% CI of the scores. **(B)** Visualizations of spatial locations of different cell populations, including all cell types (the first panel; colored by cell type annotation obtained from CellSNAP clusters; black regions indicating empty spaces) and different macrophage subpopulations identified by CellSNAP representation clusters (the second to the fourth panels). In each of the second to the fourth panels, all tumor cells, and macrophage cells from a specific CellSNAP cluster are colored, while other cells and empty spaces are in black. **(C)** Volcano plot of differentially expressed genes between CellSNAP-c6 cluster and other macrophage clusters. **(D)** Comparison of module score values (43) between CellSNAP-c6 and all other macrophage cells. ‘M1-like’ and ‘M2-like’ scores were calculated by genes from (44). Splenic macrophage specific ‘pro-inflammatory’ and ‘immunoregulatory’ scores were calculated by genes from (45). The unpaired Wilcoxon test was implemented to produce p values. **(E)** Visualization of the spatial distribution of all macrophages and their respective ligand-receptor interaction detection score levels. Detection score was calculated based on significant ligand-receptor interaction pairs between macrophages and tumor cells (46). **(F)** Top 10 most frequent ligand-receptor interaction pairs associated with CellSNAP-c6 macrophages. **(G)** GEP usage scores (47) among tumor cells, stratified by infiltration (by macrophage) status. The unpaired Wilcoxon test was implemented to produce p values.

### Application to cHL cancer tissue CODEX data

We next evaluated the performance of CellSNAP in a real-world disease context, specifically to studying the classic Hodgkin Lymphoma (cHL) tumor microenvironment (TME) and its milieu of immune infiltrates. We applied CellSNAP on 143,730 cells from an in-house 45-plex CODEX classic Hodgkin Lymphoma (cHL) tumor dataset (39). We first calculated quantitative metrics on the cell population delineation performances of the CellSNAP representation and four other representation methods (**Fig. 3A**), and implemented the cell type annotation process on either the CellSNAP representation or the feature-only representation (**Fig. 3B, Supp. Fig. 5A**). We previously identified systemic T cell dysfunction within the cHL TME using an iterative spatial proteomics and cell type-specific spatial transcriptomics approach, in which a T cell dysregulation state is directly related to its distance from neighboring cHL tumor cells (25). We postulated that CellSNAP would be an effective approach to facilitate the finer delineation of T cell subpopulations in the context of cHL. Clustering on the CellSNAP representation generated 6 distinct CD4 T cell subpopulations, with each of them occupying distinctive spatial regions relative to the tumor within the TME (**Fig. 3C, D**). In contrast, CD4 T cells obtained from clustering on feature-only representation did not exhibit these diverse spatial localizations and niche profiles (**Supp. Fig. 5B**). Two interesting CD4 T cell subpopulations identified by CellSNAP representation are (**Fig. 3D**): CellSNAP - c8 (top row, middle panel), a subpopulation that occupies the boundary of the cHL tumor patch, and CellSNAP - c20 (bottom row, right panel), a subpopulation that infiltrated inside the cHL tumor patch. We further observed that only the tumor-infiltrated CD4 T cell subpopulation (CellSNAP - c20), but not the tumor-boundary CD4 T cell subpopulation (CellSNAP - c8), exhibited an elevated expression of T cell dysfunctional markers LAG3 and PD-1 (**Supp. Fig. 5C**). These results further elucidate the importance of understanding T cell dysregulation within the cHL TME (40– 42). More importantly, the CellSNAP-enabled analysis, alongside other recent spatial-omics studies (25), shed light on the importance of spatial aspects in dysfunctional T cell - cHL TME interactions in the orchestrated immuno-logical responses to tumor in cHL and beyond.

To gain a deeper understanding of the differences between CellSNAP - c6 and other macrophages at a pathway/gene program level, we performed module scoring (43) on pan-macrophage (44) or liver-specific-macrophage public gene lists (45) (**Fig. 4D**). We found that CellSNAP - c6 macrophages skewed towards a more ‘M1-like’ and pro-inflammatory phenotype, while the other macrophages displayed a more immuno-regulatory phenotype. Given the close spatial proximity of CellSNAP c6 macrophages to specific tumor regions, and their pro-inflammatory characteristics, we hypothesized an active interaction between these macrophages and their adjacent tumor cells. To test this hypothesis, we performed spatial ligand-receptor (LR) analysis using spatialDM (46) on all macrophages and tumor cells in this tissue (**Fig. 4E**), and detected a high number of significant LR interaction events specifically enriched at the Cell-SNAP - c6 macrophage-infiltrated tumor boundary. We further performed DE analysis on LR pairs between Cell-SNAP - c6 and other macrophages, to further support a model in which the CellSNAP - c6 macrophages had increased LR interaction profiles (p.adj < 0.05, BH correction), including integrin receptors and their respective ligands compared to other macrophages (**Fig. 4F**). Our results with CellSNAP thus far identified CellSNAP - c6 as tumor-boundary infiltrating macrophages exhibiting alternative functional states and LR interaction profiles, along with the ability to more granularly distinguish HCC tumor cell populations (**Supp. Fig. 7**). We next postulated that the HCC tumor subpopulations infiltrated by these CellSNAP - c6 macrophages (CellSNAP - c7, c8, c17) would also possess different gene programs and pathway usages compared to other tumor populations even in the same tissue. To test this, we performed unsupervised Gene Expression Program (GEP) identification using cNMF (47) (**Supp. Fig. 8A, B**), and selected the most significantly upregulated or downregulated GEPs for comparison between tumor clusters with macrophage infiltrates (CellSNAP - c7, c8, c17), compared to other tumor cells in the same tissue. We then annotated the GEPs by their top 20 contributing genes, using Gene Ontology (GO) analysis (**Fig. 4G**), and observed elevated gene program activation and usages related to immune cell chemotaxis, cytokine signaling, and ECM disassembly in macrophage-infiltrated tumor cells. In contrast, other tumor cells showed enhanced gene program usages related to NOS synthesis, IL-10 regulation, and proliferation. In summary, we demonstrated that CellSNAP representation facilitated the delineation and identification of a Tumor-Associated-Macrophage (TAM) subpopulation with infiltration tendencies and pro-inflammatory functionalities. This TAM subpopulation is potentially altering tumor programs via spatial interactions, consistent with findings from previous studies in HCC tissues (49–51). Remarkably, the discovery of this TAM subpopulation is entirely unsupervised, and no training or reference dataset has been involved.

### Additional benchmarking

To evaluate the robustness of CellSNAP with respect to different tuning parameter choices, we tested ranges of values for four different tuning parameters on the mouse spleen CODEX dataset (**Supp. Fig. 9**): 1) resolution used in Leiden clustering for acquiring the cell identity cluster numbers and cell neighborhood composition. 2) K used in searching the nearest neighborhood for cell neighborhood composition calculation. 3) Image size (pixel numbers) for acquiring SNAP-CNN encoding. 4) Binarization threshold value for acquiring SNAP-CNN encoding. In addition, to evaluate the advantage of the SNAP-GNN-duo architecture, we benchmarked the model training performance, using loss values calculated on randomly selected test datasets as a proxy, of the full SNAP-GNN-duo pipeline against the single-GNN alternatives (i.e., using only the expression GNN or the spatial GNN with an MLP head) (**Supp. Fig. 10**). Altogether these results highlight the robustness of CellSNAP and its generalizable performance across multiple metrics. For details see Material & Methods.

## Discussion

Spatial-omics approaches at single-cell or even sub-cellular resolution are often limited in the number of targeted features they measure, for example *∼* 30 *−* 60 in spatial proteomic studies, and hundreds to thousands in spatial transcriptomics studies. Current computation methods for grouping biologically distinct cells in spatialomics data are often built upon those originally designed for dissociated single-cell analysis, and thus miss the opportunity to leverage the available image information and the *in situ* nature of the data for better delineation and representation of individual cells’ states.

In this study, we developed a new geometric deep learning pipeline, CellSNAP, that integrates complementary information from feature expressions (including proteins or RNAs), neighborhood context, and local tissue image, to produce for each cell a comprehensive representation vector suitable for a wide range of downstream analyses, including but not limited to clustering analysis for spatial- and-tissue-context-aware cell population delineation and identification, and analysis of their functional differences.

The CellSNAP pipeline employs a novel architecture that consists of two GNNs for coding expression similarity and spatial proximity among cells in parallel, thus allowing smooth information diffusion of biomarker expressions and local tissue images within their respective natural domains for better information fusion. CellSNAP exhibits robustness in enhancing cell population delineation across diverse spatial-omics datasets collected from different tissues and disease settings using diverse technologies. Notably, CellSNAP’s compatibility with the current *de novo* cell type identification processes allows for straightforward applications of existing clustering algorithms to the learned representation. Thus, we anticipate the adoption of CellSNAP within the spatial-omics community.

We showcased the application of CellSNAP on imaging-based spatial modalities (CODEX and CosMx-SMI). However, the pipeline would be also compatible with sequencing-based spatial modalities (e.g., Slide-seq, Stereo-seq, HDST (52–54)), as such datasets are usually accompanied by H&E or fluorescent images of the same or an adjacent tissue section, on top of the spatially-resolved genomic readouts, and hence inputs to the CellSNAP pipeline can be curated.

While the current study has focused on showcasing CellSNAP’s efficacy in cell population delineation, the utility of the learned representation vectors can be generalized to other biological tasks. For instance, they can serve as engineered features in diagonal integration tasks, complementing other recently developed methodologies (26, 27). In addition, they can serve as inputs to spatial neighborhood analysis pipelines (55, 56) for identifying different tumor-immune micro-environments and other neighborhoods of biological interests. Furthermore, the representation vectors can serve as inputs to machine learning models that aim at predicting disease outcomes directly from single-cell information. Overall, the learned all-encompassing cell representation that integrates expression, location, and image domains, summarizes key information provided by spatial-omics datasets and holds great promise for improved and better-informed down-stream analyses with diverse biological objectives.

## Materials & Methods

### CellSNAP input preparation

CellSNAP integrates single-cell feature expressions (e.g., protein or mRNA), cell neighborhood context, and local tissue image information for cell population differentiation and discovery. We first describe how to curate inputs to the CellSNAP pipeline from a typical spatial omics dataset with single-cell or sub-cellular resolution.

Assume there are *n* cells in the dataset and we have the following information. First, we have a cell-by-feature matrix *X ∈* ℝ^*n×d*^ recording *d* biomarkers (e.g., protein, gene expression, etc.) for each cell. The *i*th row of *X, x*_*i*_ *∈* ℝ^*d*^, records these biomarkers for the *i*th cell. In addition, we have spatial locations of the cells. When all cells are within the same field of view (FOV), the spatial locations can be represented as a *n×* 2 matrix storing the *x*-*y* coordinates of cell centroids within the FOV. When the dataset encompasses multiple FOVs, a FOV identifier is also recorded for each cell. Furthermore, we assume the availability of FOV(s) as digital image(s) with *C* channels, which include at least a nucleus channel and a membrane channel, similar to the set up for most imaging-based spatialomic datasets (12).

### Spatial proximity graph and feature similarity graph

We construct two graphs that share the same set of nodes, and each node corresponds to a cell in the dataset.

The first graph is a spatial proximity graph among cells. In this graph, each cell is connected to *k*_s_ spatial-nearest-neighbors within the same FOV, based on spatial coordinates of cells. We denote this graph by *G*_s_ and its adjacency matrix *A*_s_.

The second graph is a feature similarity graph among cells. For its construction, we first perform principal component analysis (PCA) on the feature matrix *X* to obtain a reduced-dimension representation 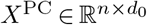. Denote its *i*th row by 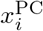 for *i* = 1,…, *n*. Then we calculate pairwise similarity of cells by Pearson correlations (or some other similarity measure that the user chooses) between row pairs of *X*^PC^ and form a *k*_f_-nearest-neighbor graph (57) by connecting each cell to *k*_f_ cells whose expression profiles are most similar. Denote this graph by *G*_f_ and its adjacency matrix by *A*_f_.

### Cellular neighborhood composition

We partition all cells into disjoint clusters by applying some graph clustering method on the feature similarity graph *G*_f_. If human-expert cell type annotations are available, we can regard them as cluster labels and skip the clustering step. For all results reported in this study, we used Leiden clustering (16) with resolution level 0.5 on *G*_f_ without human intervention for initial cell cluster label generation. These initial labels were in turn used for cellular neighborhood composition calculations.

Suppose there are in total *p* different initial cell cluster labels. Fix a positive neighborhood size *k*_1_. For the *i*th cell, its cellular neighborhood composition vector is *y*_*i*_ *∈* ℝ^*p*^ which records the proportions of its *k*_1_ spatial-nearest-neighbors belonging to each of the *p* clusters (19, 58). The spatial-nearest-neighbors of each cell are determined by the spatial coordinates of cells within the same FOV. By definition, the elements of *y*_*i*_ are non-negative and sum to one. We stack the *y*_*i*_’s as row vectors to form *Y ∈* ℝ^*n×p*^.

The generated fixed-dimensional cellular neighborhood composition vector for each cell will then be concatenated with each cell’s one-hot cluster label vector. The stacked 2*p*-dimensional vectors 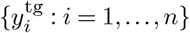 will be used as the self-supervising target of prediction in the later SNAP-GNN-duo model.

### Local tissue image tensor

For the *i*th cell, we crop its corresponding FOV at each channel *c* of interest, centered at the centroid of the cell and with window size *L × L*, resulting in a matrix 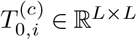. We then dichotomize the cropped image at each channel *c* at the *α*th quantile *s*_*c*_ of all values in 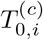 to obtain an *L× L* matrix whose (*a, b*)th entry is

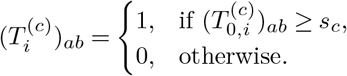

We concatenate the cropped and dichotomized images for all selected channels centered at each cell to generate the final output of this step: a 0-1-valued local tissue image tensor *T*_*i*_ *∈* ℝ^*C×L×L*^ for each cell *i ∈ {*1, 2 …, *n}*, where *C* is the total number of image channels used.

### Default tuning parameter choices for input curation

By default, we set *d*_0_ = 25 and *k*_f_ = 15 for constructing the feature similarity graph, and *k*_s_ = 15 for the spatial proximity graph. For initial cell clustering, we apply Leiden clustering with resolution 0.5. In addition, we set *k*_1_ = 20 when computing cellular neighborhood composition vectors. For image processing, the default choices are *α* = 0.9, *L* = 512 (corresponding to 50 *∼* 100*µm*), and *C* = 2 (corresponding to nucleus and membrane channels only to achieve robustness with respect to spatial omics technology).

### SNAP-CNN for encoding local tissue image

The first step of the CellSNAP pipeline trains a Convolutional Neural Network (CNN) (33), called SNAP-CNN, that predicts each cell’s neighborhood composition vector *y*_*i*_, with its local tissue image tensor *T*_*i*_. After SNAP-CNN is trained, we take the fitted hidden state of each cell as an encoding vector of the cell’s local tissue image information.

#### SNAP-CNN encoder architecture

The SNAP-CNN encoder enc(*·*) maps each local tissue image tensor *T*_*i*_ to a code *z*_*i*_ *∈* ℝ^*q*^, i.e.

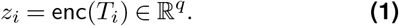

We adapt the architecture of AlexNet (59) to form a six-layer CNN as the encoder. The kernel sizes of the convolutional layers are 2, 3, 4, 2, 2, and 2, respectively, with corresponding feature map dimensions 512, 128, 63, 45, 7, 3, and 2, and channel dimensions 2, 16, 32, 64, 128, 256, and 512. The architecture also includes a 2 *×* 2 maxpooling function (33) after each of the first five convolution layers. Following the convolution layers, we flatten the feature map and add three fully connected layers with output dimensions 1024, 512, and 128, respectively. The final output dimension of the SNAP-CNN encoder is thus *q* = 128. To map *z*_*i*_ to a predicted neighborhood composition vector 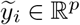, we add a fully connected layer and a non-linear activation function on top. In particular, we set

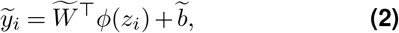

where 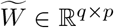 and 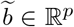 are learnable parameters and *ϕ* is the rectified linear unit (ReLU) function:

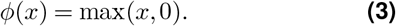

See **Supp. Fig. 1** for a schematic plot of the SNAP-CNN architecture.

After training, this step outputs fitted coding vectors *{z*_*i*_ *∈* ℝ^*q*^ : *i* = 1,…, *n}* for use in subsequent steps of CellSNAP.

#### SNAP-CNN training

The learnable parameters in enc(*·*) and in Eq. (2) are trained jointly by minimizing the mean squared error (MSE) between the true and predicted neighborhood composition vectors. In other words, the loss function for training SNAP-CNN is

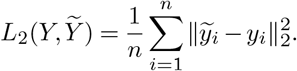

In our implementation, SNAP-CNN is trained by the Adam optimizer (60) with learning rate 10*−*4, weight decay 0, and batch size 64 for 500 epochs. To reduce overfitting and improve generalization, we apply random rotation and random horizontal and/or vertical flips on the local tissue image tensors as data augmentation during training.

### SNAP-GNN-duo for learning cell population representation

In the second step of CellSNAP, which we call SNAP-GNN-duo, we train a pair of graph neural networks (GNNs) (61) with an overarching multi-layer perceptron (MLP) head (33) to predict each cell’s neighborhood-composition-plus-cell-cluster vectors, using both its feature expressions and its local tissue image encoding. The nodes of the graphs underlying the two GNNs are shared and correspond to the individual cells in the dataset. The underlying edge structures among the nodes are different and are given by the feature similarity graph *G*_f_ and the spatial proximity graph *G*_s_, respectively. For the *i*th node (i.e., cell), its nodal covariates in *G*_f_ include feature expressions *x*_*i*_ *∈* ℝ^*d*^, its nodal covariates in *G*_s_ include local tissue image encoding *z*_*i*_ *∈* ℝ^*q*^. By training this the learnable parameters in the SNAP-GNN-duo architecture, we aim to smooth feature expression and local tissue image information based on message passing on *G*_f_ and *G*_s_, respectively, and to integrate them via the overarching MLP to predict the target vector for each cell. The cell state representation will be the concatenation of the outcomes of message passing on the pair of graphs after the model is properly trained.

#### SNAP-GNN-duo architecture

For message passing on the feature similarity graph *G*_f_, we first curate initial nodal covariate at node *i* as

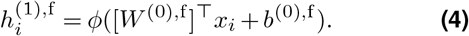

Here, 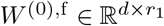 and 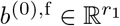 are learnable parameters, and the ReLU activation function *ϕ* (defined in Eq. (3)) is applied elementwisely on the entries of the vector in parentheses.

The collection of initial nodal covariate vectors 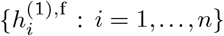 is used as the input to a GNN with network structure *G*_f_ (i.e., expression GNN in **Fig. 1**). To enable message passing across neighboring cells, we adopt the notion of graph convolution layer from (32), which can be viewed as a localized first-order approximation to spectral graph convolution (62). Stack 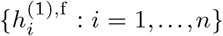 as row vectors of 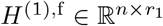. Define *Ã*_f_= *A*_f_+I_*n*_, where *A*_f_ is the adjacency matrix of *G*_f_ and *I*_*n*_ is the *n*-by-*n* identity matrix. The matrix *Ã*_f_ is the adjacency matrix of the graph 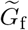 obtained from adding a self-loop to each node in *G*_f_. Further define 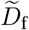 as the diagonal matrix with 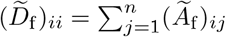, for *i* = 1,…, *n*, which records the degree of each node in 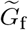. With the foregoing definitions, the message passing convolution on *G*_f_ can be written as follows:

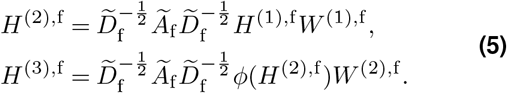

Here, 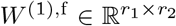 and 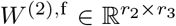 are learnable parameters. The ReLU activation function *ϕ* is applied elementwisely on *H*^(2),f^. The row vectors of *H*^(2),f^ and *H*^(3),f^ are denoted by 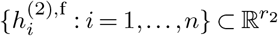 and 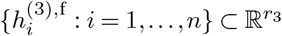, respectively.

In an analogous way, we define 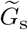, 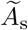, and 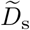 based on the spatial proximity graph *G*_s_. Stack the output of SNAP-CNN for each cell into *Z ∈* ℝ^*n×q*^. The initial nodal covariate and message passing on the graph neural network with network structure *G*_s_ (i.e., spatial GNN in **Fig. 1**) is defined as

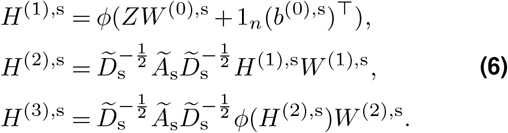

Here, 1_*n*_ is the all-one vector in 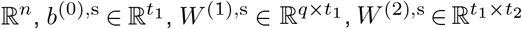, and 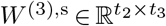 are all learnable parameters, and the ReLU function *ϕ* is applied elementwisely in both instances. The row vectors of *H*^(2),s^ and *H*^(3),s^ are denoted by 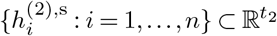 and 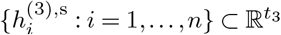, respectively.

Given the outcomes of message passing on both graphs, we define for *i* = 1,…, *n*,

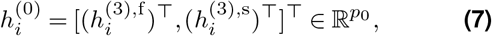

as the input to the MLP head for predicting the neighborhood-composition-plus-cell-cluster-annotation vector. Here *p*_0_ = *r*_3_ + *t*_3_. Stack 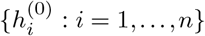 as rows of 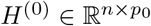. The next layers of the MLP are defined as

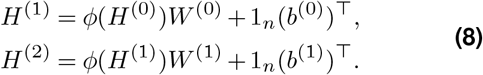

Here 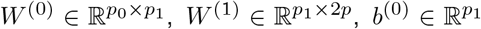, and 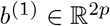 are all learnable parameters, and ReLU activation *ϕ* is applied elementwisely.

Finally, we define

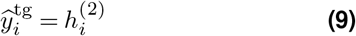

as the predicted neighborhood-composition-plus-cell-cluster-annotation vector for the *i*th cell, where 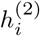 is the *i*th row vector of *H*^*(2)*^, and

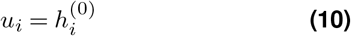

as its representation. The representation vectors *{u*_*i*_ : *i* = 1,…, *n}* are used in downstream analysis.

#### SNAP-GNN-duo training

Given the predicted and the ground-truth target vectors, i.e., 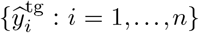 and 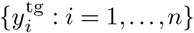, we define the loss function of SNAP-GNN-duo as

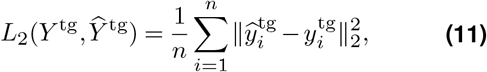

where 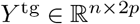 is the matrix of concatenated neighborhood composition and cell cluster vectors, and 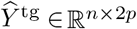 is a matrix with the *i*th row being the predicted vector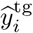 .

By default, the hidden dimensions of SNAP-GNN-duo are set at *r*_1_ = *r*_2_ = *t*_1_ = *t*_2_ = 32, *r*_3_ = 33, *t*_3_ = 11, *p*_1_ = 33. By definition *p*_0_ = *r*_3_ + *t*_3_ = 44. SNAP-GNN-duo is trained by the Adam optimizer, with a learning rate of 10*−*3 and weight decay 0, and with a single batch for 3000 epochs.

To improve the stability of the learned embedding vectors *{u*_*i*_ : *i* = 1,…, *n}*, we propose to train SNAP-GNN-duo for *m* rounds. In the *l*th round, we stack the learned embeddings as row vectors of 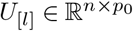. After obtaining *U*_[*l*]_, *l* = 1,…, *m*, we construct

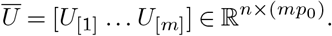

Then, we compute the *p*_0_ leading left singular vectors of *Ū*, collected as the columns of 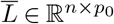, and the correspond singular values, collected as the diagonal entries of the diagonal matrix 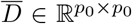. Our final learned embedding vectors are taken to be the row vectors of 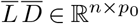, i.e.,

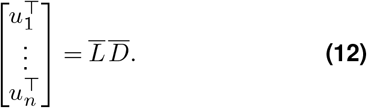

See **Supp. Fig. 1** for a schematic plot of the SNAP-GNN-duo architecture and the representation generation.

### Clustering CellSNAP representation vectors and cell population identification

The CellSNAP representation vector of the *i*th cell is 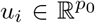 that combines complementary information from multiple domains in spatial omics data. The additional information provided by cellular neighborhood and local tissue images enables improved cell population differentiation and discovery at a finer granularity than that enabled by feature expression alone.

To this end, we construct a new graph *G*_e_ with cells as nodes. Each cell *i* is connected to its *k*_e_-nearest-neighbors measured by Euclidean distance in the representation vector space. In other words, cell *j* is connected to *i* in *G*_e_, if *u*_*j*_ is among the *k*_e_ closest points to *u*_*i*_ in 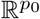, or vice versa. Then, we perform Leiden clustering (16) on *G*_e_ to find cell clusters, followed by cell-type annotation processes, similar to the procedure performed in conventional single-cell studies. For benchmarking purposes, all final clusters presented in this study used the same set of parameters for Leiden clustering (resolution = 1). Recall that each cell has a coarse initial cell population label. In the last step, we refine the clustering result on CellSNAP representation by singling out within each cluster any initial cell population that is primarily contained in the current cluster, provided that the current cluster also includes a substantial fraction of cells from other initial cell populations (by default, when the composition of initial cell populations within the current cluster, when viewed as a discrete probability distribution, leads to a Shannon entropy *>* 0.75). Further detail is documented in function `cluster_refine` in the GitHub repository.

### Evaluation and benchmarking

In this section, we define evaluation metrics for clustering, which are used in benchmarking. The objective of clustering is to minimize within-cluster variation while maintaining well-separated, distinct clusters. Consequently, compactness and separation (63, 64) serve as two key criteria for clustering evaluation.

Let

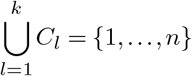

be a clustering, i.e., a disjoint partitioning, of *n* cells. Suppose we are given a feature vector 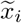 for each cell, which can be the feature expression vector *x*_*i*_ or the CellSNAP learned representation *u*_*i*_. For any vectors *x* and *x*^*′*^ of the same dimension, let

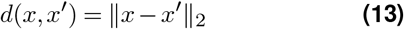

be the Euclidean distance between them.

#### Silhouette score

The Silhouette score (65) evaluates clustering performance based on the pairwise difference between within-cluster and between-cluster distances. Fix a cluster *C*_*l*_. For cell *i ∈ C*_*l*_, we first calculate

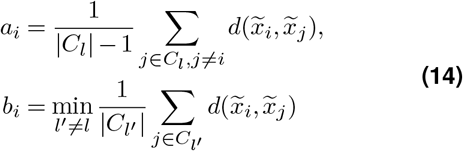

as the average within-cluster distance and the smallest average between-cluster distance for cell *i*, respectively. Here, |*C*_*l*_*′* | denotes the size of the *l*^*′*^th cluster for *l*^*′*^ = 1,…, *k*. The Silhouette score for cell *i ∈ C*_*l*_ is then defined as

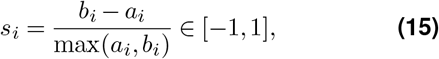

if |*C*_*l*_| *>* 1, and *s*_*i*_ = 0 otherwise. We define the Silhouette score for the clustering as 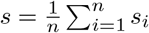. By definition, a higher Silhouette score indicates better clustering. In our benchmarking study, the Silhouette score was calculated when *K* numbers of clusters were generated by running Leiden clustering on the benchmarking representation vector, where *K* spanned from 5 *−* 30. A total of 5 batches, with each batch containing 10,000 randomly selected cells, were used for score calculation.

#### Davies–Bouldin index

The Davies-Bouldin (DB) index (66) is calculated by first computing the similarities between each cluster *C*_*l*_ and all other clusters. The highest similarity is designated as the inter-cluster separation for *C*_*l*_. The DB index is then obtained by averaging the inter-cluster separations for all clusters. Specifically, the Davies-Bouldin index for *k* clusters is defined as

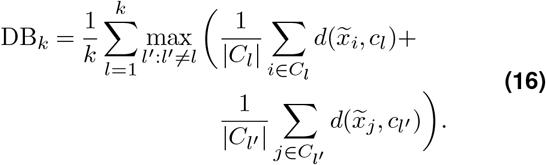

Here, for each 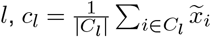 is the cluster centroid. By definition, a lower DB index value indicates better clustering performance. In our benchmarking study, the DB index was calculated when *K* numbers of clusters were generated by running Leiden clustering on the benchmarking representation vector, where *K* spanned from 5 *−* 30. A total of 5 batches, with each batch containing 10,000 randomly selected cells, were used for score calculation.

#### Calinski-Harabasz index

The Calinski-Harabasz index (CH) (67) assesses cluster compactness and separation by calculating between- and within-cluster distances. Let 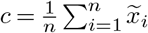 be the global data centroid. Then *d*(*c*, *c*) = *∥c − c*_*l*_*∥*_2_ is the distance between the *l*th cluster centroid and the data centroid, for *l* = 1,…, *k*. The CH index for *k* clusters is then defined as

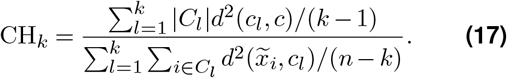

By definition, a higher CH index value indicates better clustering performance. In our benchmarking study, the CH index was calculated when *K* numbers of clusters were generated by running Leiden clustering on the benchmarking representation vector, where *K* spanned from 5 *−* 30. A total of 5 batches, with each batch containing 10,000 randomly selected cells, were used for score calculation.

#### Modularity score

When there is a graph structure among the cells, we can employ modularity (68) as a clustering validation metric. Denote by 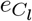 the number of edges within cluster *C*_*l*_ on the graph and 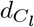 is the sum of node degrees of nodes in *C*_*l*_. Modularity score is then defined as

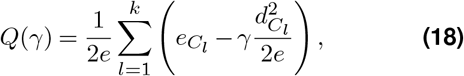

where *e* is the total number of edges. Here *γ* is a resolution parameter. In our benchmarking study, the Modularity score was calculated when different *γ* values were used. The *γ* values ranged from 0.5 *−* 2.6. A total of 5 batches, with each batch containing 10,000 randomly selected cells, were used for score calculation.

### Information retrieval efficacy evaluation of the SNAP-GNN– duo module

CellSNAP integrates information in feature expressions, neighborhood composition, and local tissue image for finer cell state differentiation. To quantitatively evaluate the additional information provided by utlizting a SNAP-GNN-duo structure in contrast to using a single GNN on either the feature or the spatial graph, we compare prediction accuracy on neighborhood composition through the following controlled experiments.

We first randomly partition all cells into a training set and a test set, with the training set containing 80% of the cells and the test set containing the remaining 20%. The overall CellSNAP pipeline was performed as described in the previous sections, with the exception: 1) when local tissue image information is excluded (feature-similarity-only-GNN), we employ the foregoing training process while omitting the SNAP-CNN step by setting *t*_1_ = *t*_2_ = *t*_3_ = 0 in the SNAP-GNN-duo architecture; 2) when feature expression profile is excluded (spatial-proximity-only-GNN), we employ the foregoing training process while omitting the feature similarity graph step by setting *r*_1_ = *r*_2_ = *r*_3_ = 0 in the SNAP-GNN-duo architecture. All other tuning parameters, the training method, and the test loss calculation remain the same.

After training, we calculate the respective loss types (L1, and L2) on the test data among the different model setups. The evaluation process was repeated 5 times, and the mean values and standard deviations were plotted.

#### Other related analysis

All detailed information related to the analysis presented in this study can be retrieved from the deposited code on GitHub, and we briefly describe them here. For all other methods benchmarked: 1) for ‘feature’, the single-cell expression profile first underwent PCA dimension reduction, and the first 25 components were used as input; 2) for ‘concat’, the 25-component feature PCA was directly concatenated with the neighborhood composition vector and used as input; 3) for ‘SpiceMix’, single-cell expression profiles along with cell spatial adjacency information were used as input for SpiceMix (31), and default parameters were implemented and trained for 200 epochs; 4) for ‘MUSE’, we first extracted images from each individual cell, using the same nuclear and membrane channels as implemented in CellSNAP. The single-cell level whole-cell segmentation masks were retrieved directly from the original data source. These images, alongside the single-cell expression profile, were used as input for MUSE (30), and the subsequent steps used default parameters. Finally, to calculate the quantitative metrics for clustering performance evaluation in each dataset across methods, a total of 5 batches, each with 10,000 randomly selected cells from the data, were used.

For the HCC CosMx-SMI data analysis, DE was performed using the R package `limma`. Module scores were calculated with the R package `Seurat` function `AddModuleScore`, with gene lists retrieved from Cheng et al. (44) or MacParland et al. (45). Spatial ligand-receptor analysis was performed with SpatialDM (46), with parameters: `l=2, n_neighbors = 30,` `single_cell=True, n_perm=200`. The LR detection score was calculated as the summation of significant (p < 0.05) LR pairs within each cell. To identify the differential LR pairs between groups, the LR detection frequency was calculated by measuring the percentage of cells with significant (p.adj < 0.05) detection of a specific LR pair, among 200 randomly selected cells, and repeated in a total of 20 batches. A Wilcoxon test (two-tailed) with Benjamini-Hochberg correction was then used to generate p-values. To define gene programs among tumor cells, cNMF, as previously described by Kotliar et al. (47), was used. The rank in cNMF (number of gene programs) was set to 25 (determined via function `k_selection_plot`). To annotate gene programs, we first selected the top 20 genes for each gene program, based on ranking from `gep_scores`, then we utilized the function `enrichr` in the R package `enrichR`, with the database `GO_Biological_Process_2015`, on these selected genes and annotated the gene programs.

All calculations and visualizations of UMAP embeddings in this study were generated using the R package `Seurat` functions `FindNeighbors` and `RunUMAP` with 30 dimensions.

## DATA AVAILABILITY

This study did not generate any new experimental data:

CODEX spleen dataset was generated from (3);

CODEX tonsil dataset was generated from (38);

CODEX cHL dataset was generated from (39);

CosMx liver dataset was generated from (48);

We have summarized all the files used in this study from the above-mentioned datasets in this link.

## CODE AVAILABILITY

CellSNAP python package, along with code used in this study, can be found in the GitHub repository: https://github.com/sggao/CellSNAP.

## ACKNOWLEDGEMENTS

The authors thank the insightful discussion with lab members from G.P.N., A.K.S., S.J., and Z.M. labs. G.P.N. is, in part, supported by the Rachford and Carlota A. Harris Endowed Professorship. A.K.S. is supported, in part, by NIH 1P01AI177687-01, Bill & Melinda Gates Foundation, Break Through Cancer, and NIH 75N93019C00071. S.J. is supported in part by NIH DP2AI171139, P01AI177687, R01AI149672, U24CA224331, a Gilead’s Research Scholars Program in Hematologic Malignancies, a Sanofi Award, the Bill & Melinda Gates Foundation INV-002704, the Dye Family Foundation, and previously by the Leukemia Lymphoma Society Career Development Program. Z.M. is supported by NSF 2345215 and NSF 2245575. S.J. and Z.G.J. are supported by BIDMC spark grant. This article reflects the views of the authors and should not be construed as representing the views or policies of the institutions that provided funding.

## AUTHOR CONTRIBUTIONS

Conceptualization: S.G., Z.M., S.C., B.Z.

Algorithm Development and Implementation: S.G., Z.M., B.Z. Analysis: B.Z., J.Y., Y.B., A.H, Y.Y.Y., G.L., S.M.

Contribution of Material and Expertise: Z.G.J., S.J.R., G.P.N., A.K.S., S.J., Z.M. Supervision: G.P.N., A.S.K., S.J., Z.M.

B.Z., S.G., and S.C. contributed equally and have the right to list their name first in their CV.

